# Engineered *Vibrio natriegens* lysate can replace multiple components of cell culture media

**DOI:** 10.64898/2026.04.09.717582

**Authors:** James Dolgin, Avani Vaid, Dylan Hendrixson, Yiming Cai, Lennie K. Y. Cheung, Licheng Xu, David L. Kaplan, Nikhil U. Nair

## Abstract

Reducing the cost and environmental impact of cell culture media is an important goal for cultivated meat, the process of generating meat *in vitro* using proliferating animal cells. While prior approaches have demonstrated the use of microbial lysates to replace expensive animal-based fetal bovine serum (FBS) in media, these formulations still rely on large quantities of growth factors such as fibroblast-like growth factor 2 (FGF2). Here, we demonstrate the use of FGF2-expressing *Vibrio natriegens* to create whole-cell lysates that replace both FBS and FGF2 in cell culture media for cultivated meat applications. This medium, named “VN40^FGF”^, supports rapid proliferation of immortalized bovine muscle satellite cells (iBSCs) in the absence of supplemented FGF2. Cells grown in VN40^FGF^ maintain phenotype and differentiation capacity. We also demonstrate that engineered *V. natriegens* can grow in spent cell culture media, further improving sustainability and economics, and reducing potential eutrophication concerns associated with waste disposal. Our approach combines multiple strategies for reducing the total number of media inputs, demonstrating opportunities for more economical and sustainable cell culture, especially for cultivated meats.

## Introduction

Cultivated meat, the process of generating meat using *in vitro* culture of animal cells, is a promising alternative to conventional livestock agriculture. Benefits of cultivated meat include streamlining food production, increasing protein availability, improving food safety, and reducing the negative externalities associated with livestock.^1,2^ However, cultivated meat has not reached significant market penetration due to its high production costs. Raw ingredient costs remain a significant roadblock to making cultivated meat reasonably accessible and affordable.

Cell culture media represents the most expensive input ingredient in cultivated meat production, comprising 98 % of raw material costs at scale.^3^ Recombinant growth factors are the primary driver of media costs, with some estimates attributing 95 % of serum-free media cost to growth factors alone.^4^ Eliminating the costly animal-derived fetal bovine serum (FBS) ingredient from cell culture media, while an important goal for cultivated meat, leads to manyfold increases in growth factor requirements.^5^ For example, immortalized bovine satellite muscle cells (iBSCs) require supplementation with numerous growth factors, including neuregulin, transforming growth factor (TGF), and fibroblast growth factor 2 (FGF2) in order to grow in the absence of FBS.^6^ These growth factors cost thousands of USD per milligram and thus impart significant costs to serum-free media (SFM). FGF2 alone, which is required at particularly high concentrations in SFM, is responsible for more than 50 % of BSC media costs.^7^ Growth factor production also carries significant negative environmental impacts, such as increases to global warming potential, ecotoxicity, and eutrophication risks.^8^ The costs of growth factors, both economic and environmental, currently serve as a significant obstacle to the low-cost sustainable production of cultivated meat.

Growth factor costs are particularly high due to their labor intensive and low-yield production process, in which they are expressed as recombinant products, typically in *E. coli* or mammalian hosts, and purified by affinity chromatography.^9,7^ Efforts to reduce growth factor costs have primarily focused on increasing yields through methods such as enhanced expression (via fusion tags, strain engineering, and metabolic optimization),^10^ bioprocess improvement,^11^ and optimization of purification.^12^ However, these innovations only marginally reduce growth factor costs. The consumables and labor associated with growth factor purification account for around 94 % of final costs.^9^ Growth factors have also been produced in spent mammalian cell culture media, but only used from these processes in their purified form.^13,14^ To avoid using purified growth factors in SFM, whole-cell lysates from growth factor-expressing microbes can be used instead. While this has been attempted previously using *Saccharomyces cerevisiae* yeast expressing a combination of four cytokines,^15^ it was not used with SFM. Our group’s previous work successfully formulated SFM – substituting recombinant albumin with lysates from the rapidly-growing bacterium, *Vibrio natriegens*.^16^ In the work, we demonstrated that *V. natriegens* lysates stimulated the growth of iBSCs at rates superior to that of a more expensive previously established Beefy-9 SFM.^6^ This established a low-cost easy to produce medium called “VN40” for its use of *V. natriegens* lysate at 40 μg protein·mL^−1^. The VN40 medium was advantageous in that it did not require large amounts of expensive purified recombinant protein, as in Beefy-9 and other studies.^6,9,17-18^ The lysate required only steps of cell growth, lysis, and filtration, making it easier to prepare than other SFM which require laborious protein extraction from plant matter or relatively slow-growing algae.^19−21^ VN is the fastest growing organism known, with a doubling time of about 10 min,^22^ and stimulates BSC growth at a low concentration of 40 μg·mL^−1^, making overall production inexpensive and rapid. However, like other cultivated meat SFM, the VN40 media still requires a relatively high concentration of supplemental commercial FGF2 of human origin. At a concentration of 40 ng·mL^−1^, FGF2 adds $ 36 L^−1^ to VN40 media costs (laboratory-scale). Fortunately, VN is a genetically tractable organism with established use in recombinant protein production.^23^ Recently, *V. natriegens* NC7 strain was developed for low resource-intensity recombinant expression, demonstrating a novel economical and highly efficient transformation method.^24^ Thus, we endeavored to engineer the *V. natriegens* NC7 strain to generate a lysate-based SFM that could be used to replace growth factors for cultivated meat production.

In this work, we demonstrate the production of a *V. natriegens* lysate containing expressed bovine FGF2 and successfully replaced both serum and FGF2 in BSC medium. This led to the development of VN40^FGF^, a serum-free medium, containing no additional FGF2. VN40^FGF^ showed comparable cell growth and phenotypic outcomes to previously developed VN40 medium, which was supplemented with human FGF2 at 40 ng·mL^−1^. We further show that the lysate can be prepared from spent mammalian cell culture media, demonstrating a closed-loop production process to further reduce production costs. In all, this study demonstrates a significant cost reduction for the process of generating cultivated meat, unlocking potential further investigations into using whole-cell lysates engineered to replace other expensive growth factors and cytokines. Overall, these findings prompt advances in cultivated meat towards an affordable reality for the general population.

## Materials & Methods

### V. natriegens transformation

A single colony of *Vibrio natriegens* NC7 (Addgene bacterial strain #215356) was used to inoculate 20 mL Minimal Competence Medium (MCM, 9 mM HEPES pH 7.4, 2 mM sodium acetate, salts). The culture was incubated at 30 °C, static, for 20 h. 25 ng of plasmid DNA, either empty vector or containing the bovine FGF2 (bFGF2) insert, was added to 350 μL culture, then incubated in static culture for 30 min (**Figure S1**). The transformation mixture was then plated directly on Lysogeny Broth (LB) + 2 % NaCl + 1.5 % agar + 2 μg·mL^− 1^chloramphenicol, then incubated at 37 °C for 1−2 days until colonies appeared.

### Extraction of microbial lysate

A single colony of *Vibrio natriegens* NC7 transformed with the bFGF2 plasmid was used to inoculate 5 mL LB + 2% NaCl + 2 μg·mL^−1^ chloramphenicol. The culture was incubated overnight at 30 °C, shaking at 200 rpm. The overnight culture was diluted 1:100 into 50 mL LB + 2 % NaCl + 2 μg·mL^−1^ chloramphenicol, then grown at 30 °C until the OD^600^ reached 0.6− 0.9. At this point, 50 μL IPTG (isopropyl β-D-1-thiogalactopyranoside) was added to the culture, and incubation continued for 4−6 h. The cell pellet was harvested by centrifuging the culture at 5,000 × g for 30 min, washing and resuspending in 15 mL of sterile phosphate buffer saline (PBS, 10 mM phosphate, 140 mM NaCl, 3 mM KCl, pH 7.4). To generate lysate, the cell suspension was sonicated (Branson 150 sonicator and 10 s Pulse, 30 s gap between each pulse and a total 7 min at 45 % amplitude) on ice. After sonication, the soluble lysate was obtained by centrifuging the sonicated solution at 21,000 × g for 20 min. The supernatant was then filtered through a 0.2 μm syringe filter (Corning #431219). The total protein content of the lysate was determined using Bradford Assay (Coomassie Plus™ Protein Assay Reagent, ThermoFisher #23238), using bovine serum albumin (BSA) to generate a standard curve. The concentration of FGF2 in the lysate was determined using the Invitrogen Bovine/Human bFGF ELISA Kit (ThermoFisher #EB2RB).

### iBSC culture and growth studies

iBSCs were previously isolated from the semitendinosus of a 5-week-old Simmental calf at the Tufts Cummings School of Veterinary Medicine.^6^ These cells were engineered to constitutively express bovine telomerase reverse transcriptase (TERT) and cyclin-dependent kinase 4 (CDK4) in prior studies.^25^ After adaptation to VN40 media as described previously,^16^ iBSCs at passage 70 were seeded in triplicate at 2,000 cells·cm^−2^ in serum-free, FGF2-free B8 medium with 1.5 μg·cm^−2^ vitronectin (rhVTN-N; ThermoFisher #A31804) in tissue culture polystyrene 6-well plates. B8 medium was formulated as previously described,^6^ omitting FGF2. After allowing iBSC adherence for at least 3 h, cells were washed with DPBS (Dulbecco’s phosphate buffered saline; ThermoFisher #14040117) and fed B8 supplemented either human FGF2 (hFGF2) at 40 ng·mL^−1^ (PeproTech #100-18B, Rocky Hill, NJ, USA) or 40 μg·mL^−1^ lysate from bFGF2-expressing *V. natriegens*. Negative controls were fed B8 with no added FGF2. Cells were fed every 2−3 days until they reached 80−90 % confluence. At this point, cells were passaged using TrypLE Express enzyme (ThermoFisher #12604013) and counted using a NucleoCounter NC-200 (Chemometec, Allerød, Denmark). Doubling time (*DT*) was calculated using **Equation 1**

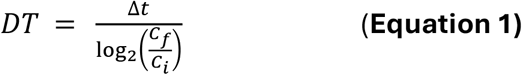

with *Δt* being time between passages, *C*_*f*_ being cell count at passage, and *C*_*i*_ being total cells seeded. At passage, cells were centrifuged at 300 × g and seeded in a new well. Cells were cultured at 37 °C and 5 % CO_2_. During feeding of iBSCs with VN40^FGF^, spent media was collected for growth of *V. natriegens* for recycled media studies. For these studies, cells were similarly fed VN40^FGF^ produced from spent media and grown and counted as described.

### Characterization and differentiation of iBSCs

After the fifth passage in VN40/VN40^FGF^ media, iBSCs were allowed to adhere in triplicate in 48-well plates at 6000 cells·cm^−2^ and their satellite cell identity was characterized via staining for Paired-box 7 (Pax7). Cells were fixed with 4 % paraformaldehyde in PBS (ThermoFisher #AAJ61899AK) for 20 min, washed with DPBS containing 0.1 % Tween-20 (ThermoFisher, BP337), permeabilized for 10 min with DPBS containing 0.1 % Triton X-100 (ThermoFisher, BP151), washed again with DPBS + Tween-20, and blocked for 30 min using a blocking solution containing DPBS with 5 % goat serum (ThermoFisher #16210064) and 0.05 % sodium azide (Sigma #S2002). A primary antibody solution containing 1:500 Pax7 antibody (ThermoFisher #PA5-68506) and 1:1000 actin stain Phalloidin 488 (ThermoFisher A12379) in blocking solution was added to cells and incubated overnight at 4 °C. Cells were then washed with DPBS + Tween-20 and blocked for 30 min once again. A solution containing 1:500 Pax7 secondary antibody (ThermoFisher #A11072) and 1:1000 nuclear stain DAPI (ThermoFisher #D1306) in blocking solution was added to cells and incubated for 1 h at room temperature. Cells were then washed with DPBS + Tween-20 and kept in DPBS for imaging. Imaging was carried out using fluorescence microscopy (KEYENCE, BZ-X700, Osaka, Japan). Stemness was quantified using a Celigo automated image cytometer (Revvity, Waltham, MA) to determine cells positive for Pax7. Cells were differentiated by seeding at 5,000 cells·cm^−2^ in 48-well plates and initiating differentiation when cells appeared confluent 3 days after initial seeding. Differentiation media contained DMEM + Glutamax supplemented with 2 % horse serum (ThermoFisher #16050130), 0.5 mg·mL^−1^ recombinant human albumin, 1 × ITS-X (ThermoFisher #51400056), 0.5 mM LDN193189 (Sigma #SML0559), 1% antibiotic/antimycotic, and 2.5 μg·mL^−1^ puromycin. Differentiation continued for 9 days with cells fed fresh differentiation media every 2−3 days. After mature myotubes were observed, cells were fixed, permeabilized, and blocked as described. Cells were then stained using myosin heavy chain (MHC; Developmental studies hybridoma bank #MF-20, Iowa City, IA, USA) and 1:200 desmin (Abcam #ab15200, Cambridge, UK) primary antibodies diluted in blocking solution. Secondary antibodies for MHC (ThermoFisher #A32723; 1:200) and desmin (ThermoFisher #A11072; 1:500) along with 1:1,000 DAPI, diluted in blocking buffer, were then applied as described. Imaging was carried out using fluorescence microscopy (KEYENCE, BZ-X700, Osaka, Japan). Differentiation was quantified using Celigo automated image cytometer to determine percentage of cells contained in desmin-positive myotubes (fusion index).

### Spent medium recycling

To generate VN40^FGF^ from spent cell culture media, we followed the scheme in **Figure 1**. The spent medium from iBSC cell culture was collected, supplemented with 3 % NaCl (w/v), and autoclaved with a 40 min liquid cycle. *V. natriegens* NC7 was cultured overnight in LB + 2 % NaCl, then subcultured into the sterilized spent media at a starting OD_600_ of 0.05. Growth curves were generated using the BioTek Epoch2 Microplate Spectrophotometer (Agilent Technologies, Santa Clara, CA, USA), taking new OD_600_ measurements every 10 min for 24 h. Spent media collected from iBSC and *V. natriegens* growth was analyzed for ammonium, glucose, glutamate, glutamine, lactate, and potassium using YSI 2950 Biochemistry Analyzer (Xylem Corp., Washington, DC), using buffer standards and sensors provided by the manufacturer.

**Figure 1.**
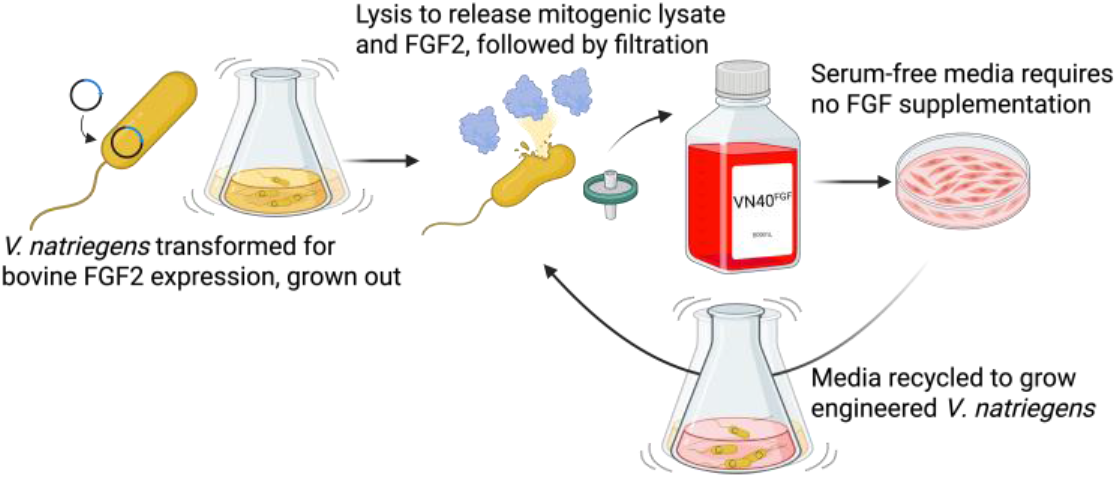
Overview of workflow. Production scheme of engineered *V. natriegens* lysates to replace commercial FGF2 using recycled spent medium.

### Statistics

Statistical analysis was carried out using GraphPad Prism v10.6. Significant measurements were carried out using Student’s t-test, or one-way ANOVA with Dunnett’s correction for multiple comparisons to detect the presence of significantly different groups.

## Results

### *V. natriegens* NC7 can be engineered to express bovine FGF2

We demonstrated the successful preparation of whole-cell lysate from *V. natriegens* NC7 strain engineered to express bovine FGF2 (bFGF2) using simple steps of growth, sonication, and filtration (**Figure 1**). Visualization of the total soluble protein content by SDS-PAGE showed the presence of a ∼17 kDa band for the recombinant *V. natriegens*, suggesting successful expression of bFGF2 relative to the parental strain (**Figure 2A**). Whole-protein concentration of the lysate was ∼3,800 μg·mL^−1^, with 66 μg·mL^−1^ (i.e., < 2 % total protein) being bFGF2 (**Figure 2B**). Thus, bFGF2 can be delivered to iBSCs at ∼700 ng·mL^−1^ when including 40 μg·mL^−1^ *V. natriegens* lysate in media,^16^ which is significantly higher than the minimum 40 ng·mL^−1^ of hFGF2 required for robust growth in serum-free conditions.^6^

**Figure 2.**
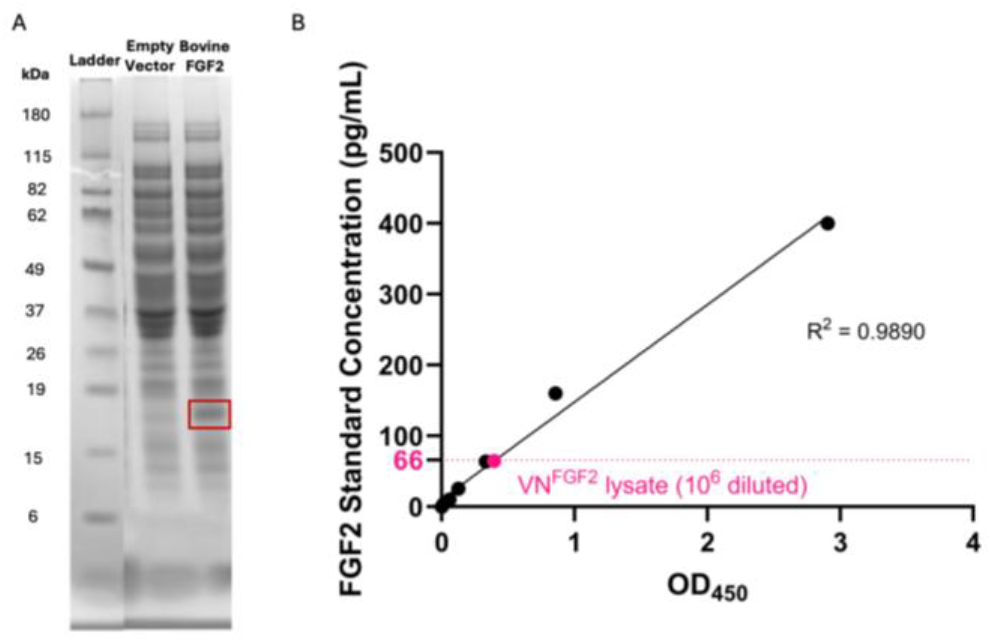
Expression of bFGF2 in transformed *V. natriegens* lysate. **A)** SDS-PAGE gel of whole-cell lysates of *V. natriegens* NC7 transformed with either the bFGF2 expression plasmid or an empty vector (EV). The band corresponding to FGF2 (∼17 kDa) is indicated. **B)** Quantification of bFGF2 concentration in *V. natriegens* lysate (10^6^-fold dilution being 66 pg·mL^−1^) using ELISA. Parental strain showed no signal in this assay. Data points of calibration curve are also shown.

### VN40^FGF^ replaces FGF2 needs in VN40 for long-term iBSC proliferation

We previously generated VN40 medium by adding lysate from wild-type *V. natriegens* at 40 μg·mL^−1^ protein and recombinant purified human FGF2 (hFGF2) at 40 ng·mL^−1^ to B8 basal medium.^16^ To create VN40^FGF^, we added lysate from bFGF2-expressing engineered *V. natriegens* at 40 μg·mL^−1^ protein to B8 medium. After adaptation to their respective media, we compared the growth kinetics of iBSCs in VN40^FGF^ and VN40 media. Cells grew at equivalent rates in both VN40^FGF^ and VN40, demonstrating the ability of expressed bFGF2 to replace supplemental hFGF2 in iBSC culture (**Figure 3A**). Doubling times were ∼32.6 h in both media (**Figure 3B**), which was slower than those previously reported for serum-containing media (∼22 h), consistent with prior studies with these cells.^16^ However, iBSCs doubled more rapidly in VN40^FGF^ than in other FGF-replacement media, such as in recent attempts to engineer iBSCs to self-express bFGF2 (66 h).^26^ We also confirmed cell morphology was comparable for cells grown in VN40^FGF^ (**Figure 3C**) and VN40 (**Figure 3D**). Thus the cells were not negatively impacted by bFGF2 expressed by *V. natriegens* or by the higher bFGF2 concentration delivered in VN40^FGF^. These results demonstrate for the first time that recombinant growth factors expressed in the fast-growing *V. natriegens* can stimulate animal cell proliferation.

**Figure 3.**
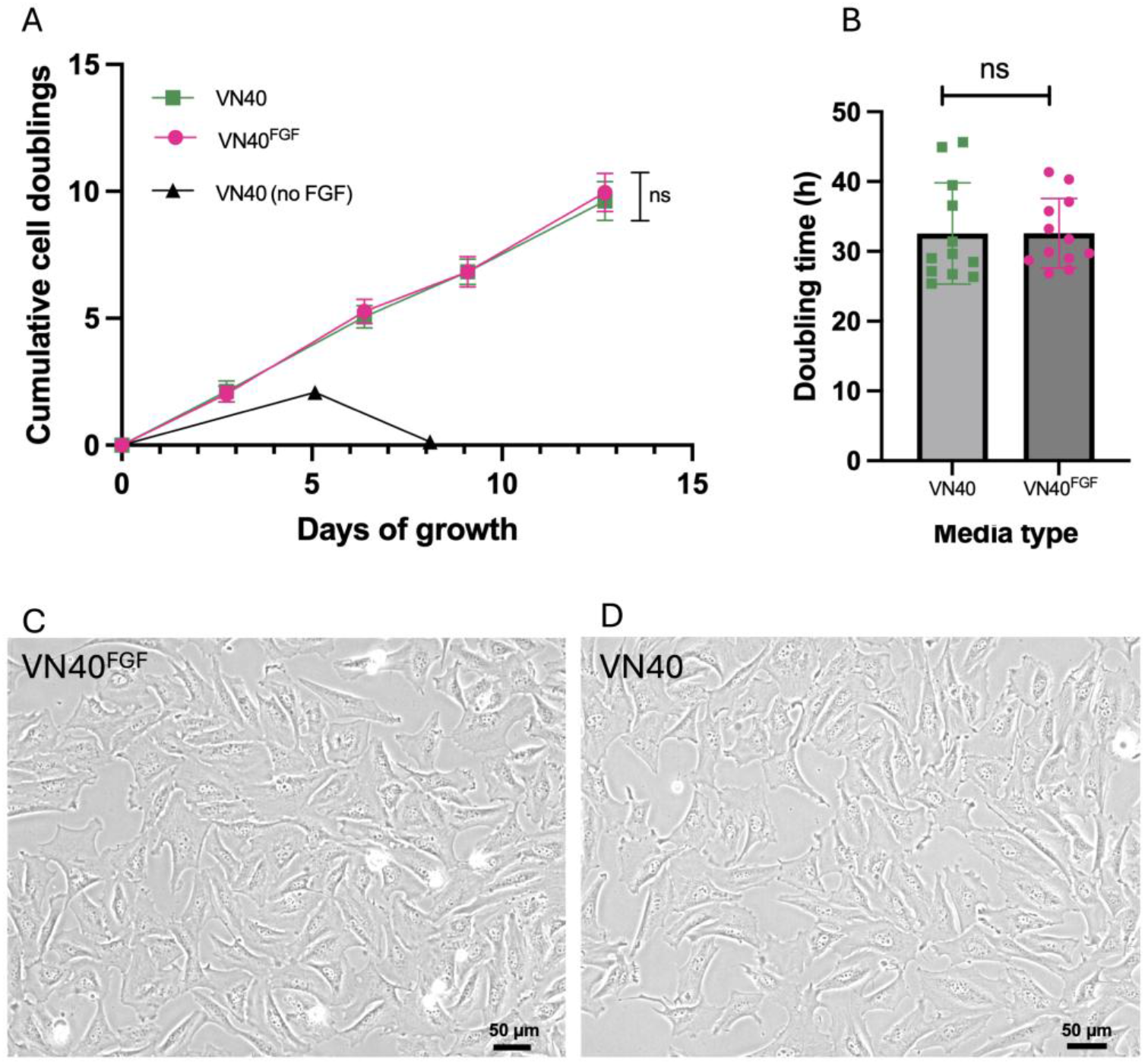
VN40^FGF^ stimulates iBSC cell growth in absence of supplemental FGF2. **A)** Multi-passage growth curves comparing iBSC growth in VN40^FGF^ medium (lysate from bFGF2-expressing *V. natriegens* in FGF-free SFM), VN40 medium (lysate from non-engineered *V. natriegens* in hFGF2-supplemented SFM), showing equivalent growth between the two, and VN40 medium with no FGF2 as negative control, showing senescence in absence of FGF2. **B)** Calculated doubling times of iBSCs growing in VN40^FGF^ and VN40, showing equivalent growth rate. p > 0.1234 (ns) by Student’s unpaired t-test. **C)** Brightfield of iBSCs after 3 days of growth in VN40^FGF^. **D)** Brightfield of iBSCs after 3 days of growth in VN40. Scale bar = 50 μm.

### iBSCs grown in VN40^FGF^ maintain stemness and differentiation capacity

Using Pax7 immunostaining, microscopy (**Figure 4A**), and quantitative image cytometry (**Figure 4B**) >98 % of the iBSCs maintained satellite cell phenotype over five passages in VN40^FGF^.^27^ This was equivalent to cells passaged in standard VN40 medium, showing bFGF2 expressed from *V. natriegens* did not negatively impact stemness phenotype in the cells. After multiple passages in VN40^FGF^, we were able to differentiate the iBSCs into mature multi-nucleated myotubes expressing Desmin and Myosin Heavy Chain proteins (**Figure 4C**), indicating early- and late-stage maturation, respectively.^28^ These results demonstrate that cells were able to differentiate successfully despite the higher FGF2 concentration from extracts of VN40^FGF^. In fact, cells grown in VN40^FGF^ differentiated with higher myotube formation than cells grown in VN40 (**Figure 4D**), perhaps due to higher concentrations of FGF2 impacting extracellular signal-regulated kinase (ERK) pathways and causing increased differentiation as shown previously.^29^

**Figure 4.**
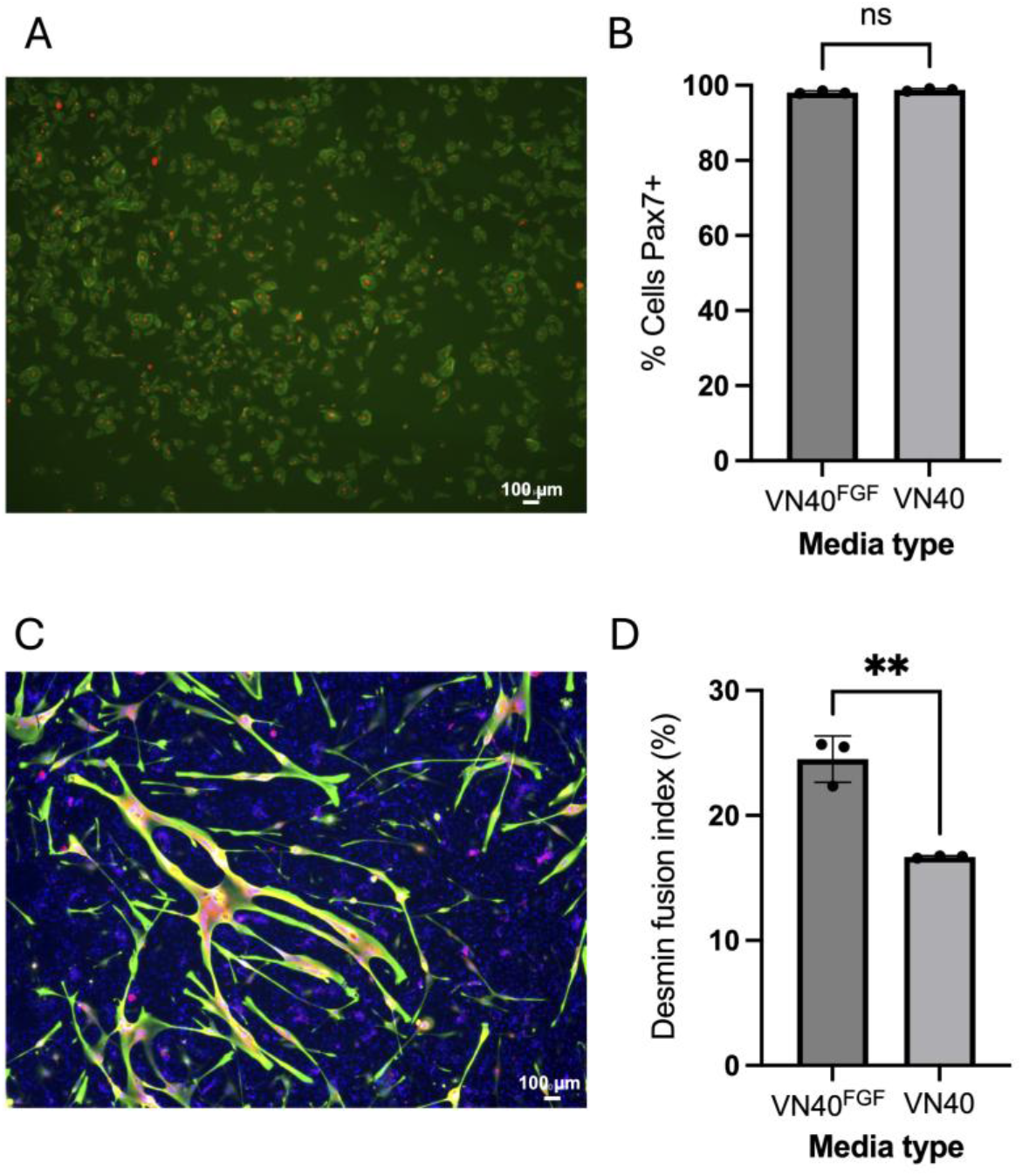
iBSCs grown in VN40^FGF^ maintain phenotype and differentiation capacity. **A)** Pax7-based immunostaining of iBSCs maintained in VN40^FGF^ for five passages. Red = Pax7 stain, Green = Phalloidin actin stain, Scale bar = 100 μm. **B)** Pax7^+^ cells according to automated image cytometry, quantifying % Pax7^+^ Phalloidin^+^ cells compared to iBSCs grown in VN40 medium. **C)** Differentiation marker immunostaining of iBSCs maintained in VN40^FGF^ for five passages and differentiated for 9 days. Red = Myosin Heavy Chain, Green = Desmin, Blue = DAPI nuclei stain, scale = 100 μm. **D)** Fusion index of differentiated iBSCs grown in VN40^FGF^ compared to VN40. Fusion determined by % DAPI^+^ cells in desmin^+^ myotubes. p > 0.1234 (ns), p < 0.0021 (**) by Student’s unpaired t-test.

### *V. natriegens* lysate can be produced from spent cell culture media

To increase sustainably and circularity of the process, we grew engineered *V. natriegens* in spent VN40^FGF^ after iBSC growth. A potential obstacle in growing *V. natriegens* in spent cell culture medium is the presence of antibiotic-antimycotic (anti-anti), which is added to mammalian cell culture medium to prevent contamination. Treating the media with a 40 min autoclave cycle enabled *V. natriegens* growth, suggesting that autoclaving destroyed the antimicrobial components in the media. To investigate the limitations of DMEM/F12 media on *V. natriegens* growth, we measured *V. natriegens* growth over 20 h in LB, fresh VN40^FGF^, and spent VN40^FGF^ (**Figure 5A**). All media conditions were supplemented with salt to mimic typical saline growth conditions of *V. natriegens*. VN40^FGF^ media, both spent and fresh, supported growth of *V. natriegens*, though rich media (LB + 2 % NaCl) yielded a greater maximum OD_600_ (1.37 for LB vs. 0.5 for spent VN40^FGF^).

**Figure 5.**
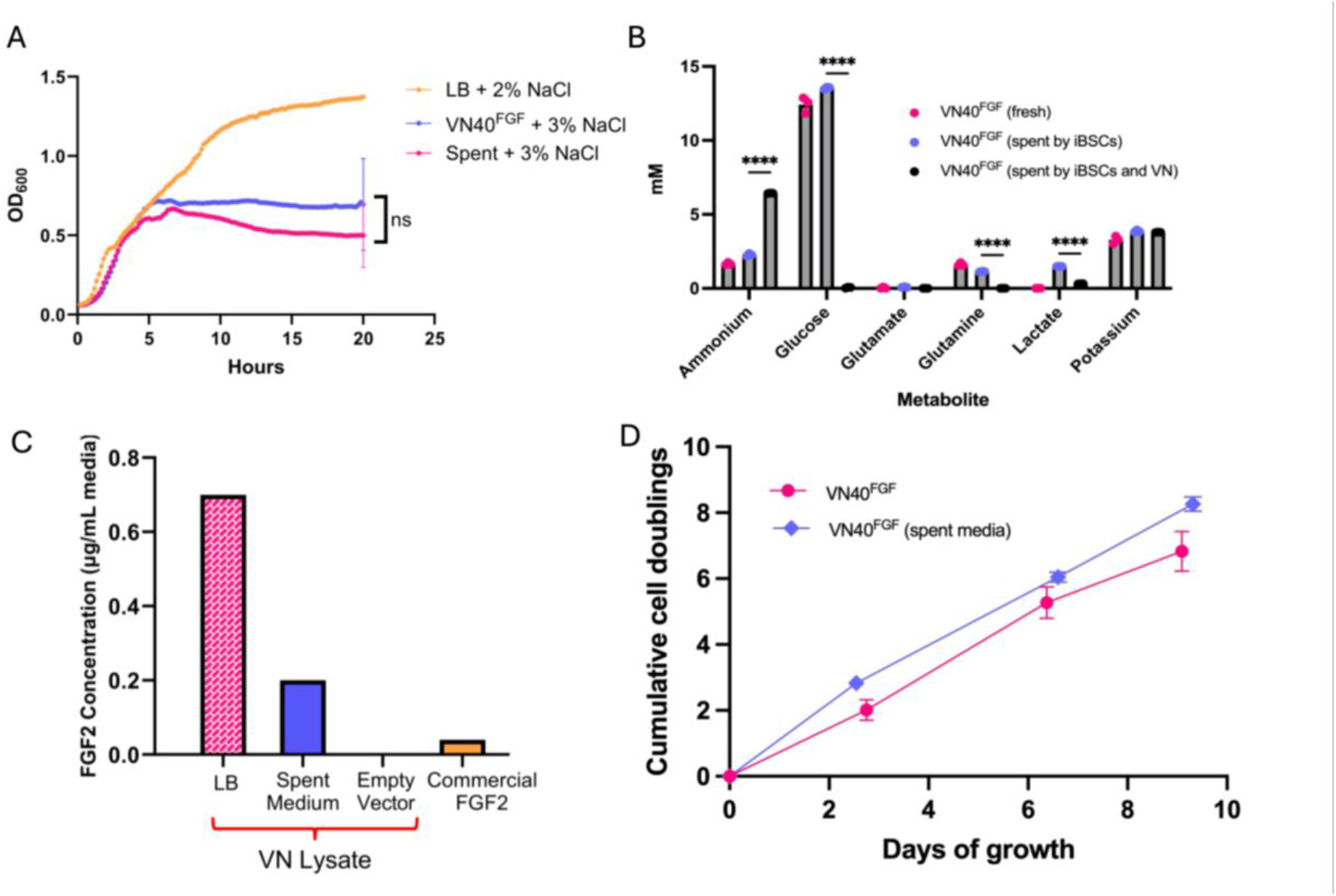
Spent medium recycling enables circular VN40^FGF^ lysate production. **A)** *V. natriegens* growth in salt-supplemented and autoclaved LB, fresh VN40^FGF^, and spent VN40^FGF^. One-way ANOVA with Dunnett’s correction for multiple comparisons, p < 0.05 (ns). Data points shown are the mean of three replicates. **B)** Metabolite analysis of fresh VN40^FGF^, spent VN40^FGF^ after growth of iBSCs (with media collection every 2−3 days), and spent medium after growth of *V. natriegens*. One-way ANOVA with Dunnett’s correction for multiple comparisons, p < 0.0001 (****). **C)** Quantification of bFGF2 delivered in VN40^FGF^ by *V. natriegens* lysate grown in LB (700 ng·mL^−1^) and spent medium (212 ng·mL^−1^) compared to empty vector control (< detection limit) and concentration of commercial FGF2 added to SFM media. **D)** iBSC growth in VN40^FGF^ medium containing either lysate grown in LB or spent medium over 9 days (n = 3 replicates at each timepoint).

To determine nutrient composition of spent VN40^FGF^ medium, we analyzed the metabolites,^30^ and found increases in ammonium and lactate, a decrease in glutamine, and no change in glutamate, potassium and glucose (**Figure 5B**). After growing *V. natriegens* in spent media, we found almost complete consumption of glucose, glutamine, and lactate and significant production of ammonium. Thus, *V. natriegens* rapidly consumes the organic components of spent culture medium (likely glucose and glutamine first,^31^ followed by lactate).^32^ Spent VN40^FGF^ is therefore sufficiently nutrient-rich to support its use as recycled medium for *V. natriegens* growth, though growth could be improved through supplementing spent media with additional substrates. When growing *V. natriegens* in LB, we similarly observed ammonium accumulation (**Figure S2**), showing growth in both media are limited by the availability of carbon, not nitrogen.

The amount of bFGF2 present in *V. natriegens* lysate was quantified using ELISA. *V. natriegens* grown in spent medium yielded less bFGF2 than that grown in LB (212 vs. 700 ng·mL^−1^ in VN40^FGF^), which was consistent with their respective growth profile (**Figure 5C**). However, both media conditions generated a large excess of bFGF2 compared to the amount of commercial FGF2 that is added to SFM (40 μg·mL^−1^). Using VN40^FGF^ generated from spent media resulted in iBSC growth similar to VN40^FGF^ generated from standard LB (**Figure 5D**), suggesting the activity of the lysate and recombinant growth factor is not negatively impacted by lower expression in the spent growth medium. We show that spent VN40^FGF^ medium pooled from three passages of iBSCs was sufficient to re-generate VN40^FGF^ for three additional passages. This shows an efficient media recycling process in which LB inputs can be minimized during manufacturing.

### VN40^FGF^ produced using spent medium is cost-competitive in comparison to previous iBSC growth medium formulations

To determine the cost-benefit of replacing commercial hFGF2 with our engineered lysate, we performed a analysis by tabulating the reagent costs of each media formulation iteration (**Figure 6**). The previous VN40 formulation had resulted in a significant cost reduction with respect to serum-containing bovine satellite cell growth medium (BSCGM), reducing media cost from $323 L^−1^ to $158 L^−1^, a 51 % reduction (**Table S1−S3**).^16^ By replacing hFGF2 in the media with whole-cell lysate, we further lower the media cost to $124 L^−1^, achieving a further 22 % cost reduction with respect to VN40 and a 62 % reduction with respect to BSCGM (serum-containing media) (**Table S3−S4**). Next, we determined the cost benefit of growing *V. natriegens* in recycled spent medium rather than LB. The reagent costs to produce *V. natriegens* lysate using LB equate to $4.50 L^−1^ of VN40, but by growing the bacterium in spent medium, we reduce this cost by 72 % to $1.25 L^−1^ **(Table S5−S6)**. Importantly, these costs represent reagents purchased for lab-scale production. Media costs are expected to fall several orders of magnitude with large-scale amino acid production, shifts away from pharmaceutical-grade materials, and further decreases to growth factor costs.^33^

**Fig. 6.**
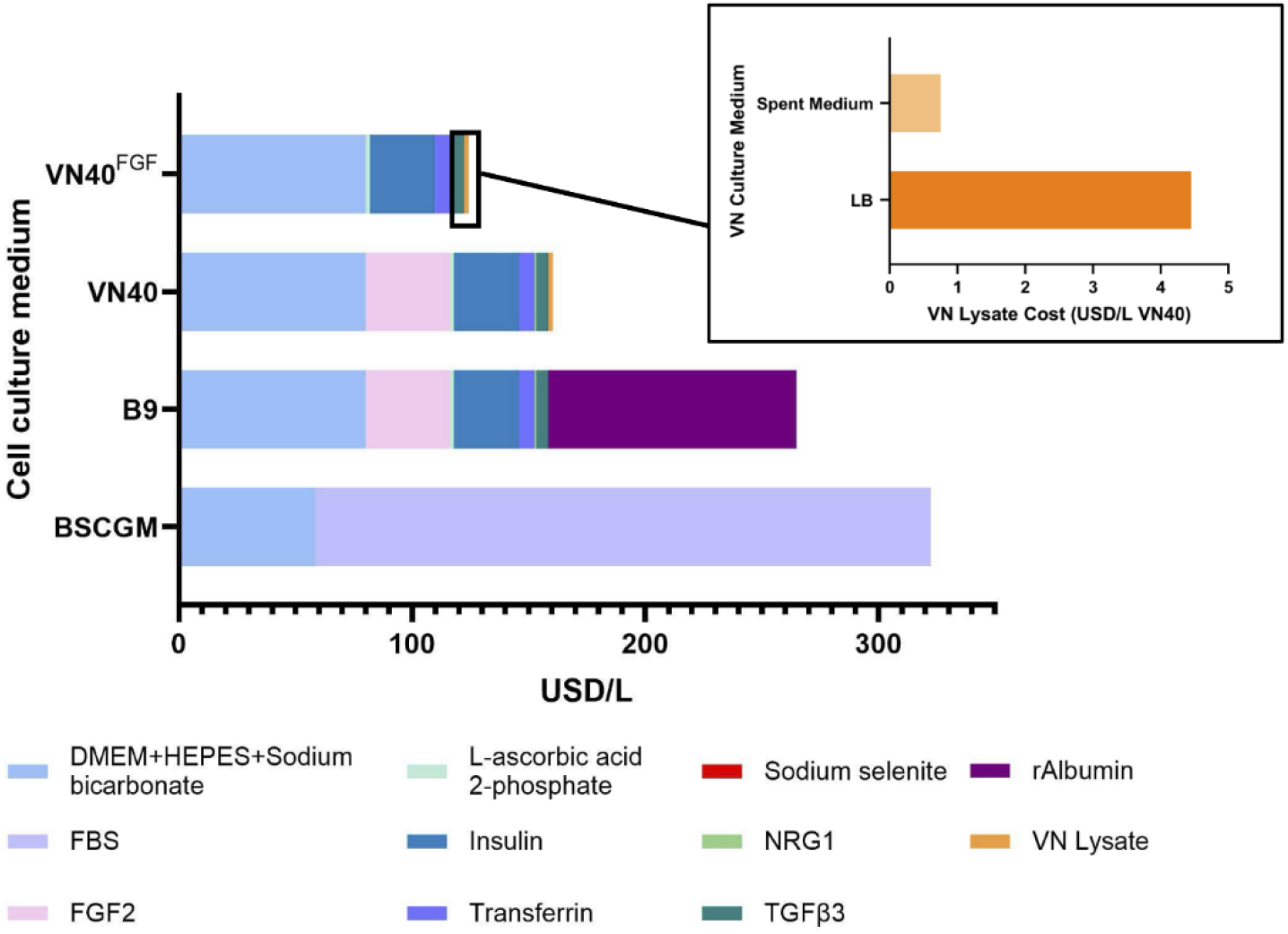
Media cost analysis. Cell culture media costs for bovine satellite cell growth medium, Beefy-9, VN40, and VN40^FGF^, assessed based on the cost of each component in USD·L^−1^ medium.

## Discussion

Cultivated meat offers a potential solution to mitigate the environmental impact of livestock generated meat for consumption. However, this process must be cost-effective and sustainable to achieve widespread accessibility and appeal. The resource burden of cell culture media is a significant roadblock to this goal.^3,34^ The high cost and environmental impact of media for cultivated meat is largely attributed to the use of fetal bovine serum (FBS) and intensive growth factor purification processes.^4,8^ Simplifying growth factor preparation can lead to significant reduction in resource needs for media production. Recently, we developed VN40 as an SFM using nutrient-rich whole-cell lysates from the rapidly dividing *V. natriegens* bacterium.^16^ This medium was advantageous, as it did not involve lengthy extraction or purification processes,^9,17,19,21,35-36^ instead requiring only a simple process of mechanical lysis and filtration to produce serum-replacing components. Still, VN40 contains purified, recombinant FGF2, which contributes significant cost. Further, the media used to grow *V. natriegens* may contribute to increased costs and environmental impact at scale.^37-38^ Our work shows that by engineering *V. natriegens* to express bFGF2, we can generate whole-cell lysates to replace both serum and hFGF2 in cultivated meat production without resource-intensive expensive purification processes. To further reduce the costs of *V. natriegens* culture, we also showed that these lysates can be produced circularly using spent media from mammalian cell culture.

*V. natriegens* has been used primarily for production of therapeutic proteins,^23^ but here we also demonstrate its production of a recombinant growth factor. Growth factors like FGF2 and insulin-like growth factor 1 (IGF-1) used for cultivated meat production are typically produced in recombinant *E. coli*.^10^ *V. natriegens* possesses several potential advantages over *E. coli* for growth factor production – such as improved protein folding,^39^ reduced endotoxin content,^40^ and rapid growth rate.^41^ Combining this bacterium’s efficient biomass production rate and recombinant expression abilities, we developed VN40^FGF^ from whole-cell lysates of bFGF2-expressing *V. natriegens*. With one liter of *V. natriegens* culture, we can produce 152 mg total lysate protein, enough for almost 4 L of cell culture media. While serum-free iBSC media, such as VN40, contains a controlled quantity of 40 ng·mL^−1^ hFGF2, the VN40^FGF^ contained about VN40^FGF^ elicited iBSC growth comparable to VN40. This implies that bFGF2 from *V. natriegens* functions properly for binding and activation of bovine FGF receptors, and that iBSCs are tolerant of higher FGF2 concentrations than previously explored.^9^ Further studies could explore the dose-dependent potency of *V. natriegens* versus *E. coli-*expressed bFGF2. Further, total bGFG2 could be titrated to lower levels if needed via dilution with extracts from the untransformed *V. natriegens*. While we used a bovine FGF2, future studies can explore proteins sourced from other species as these could prove to be more potent.^9^ The use of bovine FGF2 is advantageous over human FGF2 used in prior studies from a regulatory and consumer acceptance standpoint.^6,16^

iBSCs demonstrated maintenance of satellite cell phenotype and differentiation capacity after growth in VN40^FGF^, despite the overabundance of bFGF2 present in this medium. While FGF2 is primarily a cell cycle progressing factor, studies point to its role in stimulating muscle regeneration and differentiation,^29,42^ which could explain higher differentiation in VN40^FGF^-fed cells. Evidently, residual bFGF2 remaining after the removal of proliferation medium did not cause issues for differentiation. This is cited as an issue in studies exploring engineering bovine cells for self-signaling bFGF2 production,^26^ where its biosynthesis cannot be turned off, causing interference with differentiation. VN40^FGF^ also does not rely on engineered bovine cells, which could raise regulatory and consumer acceptance concerns. While we demonstrate that our lysate promoted iBSC growth and differentiation to an equal or greater extent as did commercial FGF2-containing media, we note that the stability and structure of our *V. natriegens*-generated bFGF2 has not been investigated. Under the cultivation conditions utilized here, it is sufficiently stable to have a positive biological impact.

We saw an additional opportunity to reduce resource burden of VN40^FGF^ production by cultivating *V. natriegens* on spent cell culture media rather than using fresh bacterial growth medium. This promises to significantly reduce overall cost and environmental impact of cultivated meat media.^43,44^ We fully repurposed our spent iBSC cell culture media for VN40^FGF^ growth, which was then used for equivalent cell growth. This demonstrates a fully circular process, where mammalian cell growth can subsist entirely on VN40^FGF^ produced from prior passages. While growth of *V. natriegens* on spent media was inferior to that in LB, sufficient lysate was produced for additional cycles of iBSC growth. *V. natriegens* growth in spent VN40^FGF^ did not differ with fresh VN40^FGF^, though analysis of spent VN40^FGF^ media revealed an increase in ammonium and lactate, and a decrease in glutamine, compared to fresh VN40^FGF^. These metabolite differences did not impact nutrient availability for *V. natriegens* to successfully grow, though higher densities of suspension mammalian cell culture may cause higher nutrient depletion. Prior spent media analyses suggest the rapid consumption of glutamine by mammalian cells, but a more delayed consumption of glucose,^45^ potentially explaining the unchanged glucose content during frequent media changes. Nutrient availability for *V. natriegens* growth showed that the cell culture media could be further recycled through glucose and glutamine supplementation, which could also induce VN ammonium consumption.^46^ Our results showed *V. natriegens* may be used for lactate removal to regenerate media for mammalian cell culture, which would recycle media and produce lysate simultaneously.

This work results in 22 % cost savings compared to VN40, and 62 % cost-savings compared to serum-containing media. Growth factors represent a significant cost in cultivated meat, and could comprise a greater share of cost at scale compared to other media ingredients.^47^ Basal media represents the next most expensive media ingredient, though these costs are expected to drop dramatically upon scale.^48^ Expression of other growth factors, such as TGFβ3 and transferrin, in engineered lysates could further decrease costs of media especially at scale. Processing improvements for lysate production, such as high-pressure homogenization instead of sonication, would further decrease costs at scale.^37^ This work also results in significant decreases in resource use for cultivated meat. Growth factor purification involves resource-intensive column separation and dialysis, the need for which this work eliminates.^8^ Further resource demands and environmental externalities are caused by the fermentation step of growth factor production.^8^ Here, we reduce these burdens by demonstrating growth of engineered *V. natriegens* in spent media. Resource demand can be further reduced by taking advantage of ambient temperature fermentation.^24^ Overall, VN40^FGF^ shows a low-cost and low-resource usage approach to creating SFM for cultivated meat that does not require supplemental growth factor addition. This work brings cultivated meat closer to reaching affordability and sustainable production.

## Acknowledgements

This study was supported by the Tufts University Center for Cellular Agriculture (TUCCA) – Consortium to N.U.N and D.L.K., and the USDA grant 2021-05678 (D.L.K.). We thank Dr. Ben Richardson (Upside Foods), Dr. Elliot Swartz (Good Food Institute), Dr. James Brooks (ThermoFisher Scientific) and Stacy Holdread (ThermoFisher Scientific) for providing technical input. We would also like to thank members of the Nair and Kaplan group for their helpful insights.

## Author Contributions

James Dolgin: Methodology, Formal analysis, Investigation, Writing - Original Draft, Review & Editing, Visualization, Avani Vaid: Methodology, Formal analysis, Investigation, Writing - Original Draft, Review & Editing, Visualization, Dylan Hendrixon: Investigation, Yiming Cai: Investigation, Lennie K. Y. Cheung: Investigation, Licheng Xu: Investigation, David L. Kaplan: Writing - Review & Editing, Supervision, Funding acquisition, Nikhil U. Nair: Conceptualization, Methodology, Writing - Review & Editing, Supervision, Funding acquisition.

## References

1 J. Cai, S. Wang, Y. Li, S. Dong, J. Liang, Y. Liu and S. Li, Industrialization progress and challenges of cultivated meat, Journal of Future Foods, 2024, 4, 119–127. 10.1016/j.jfutfo.2023.06.002.

2 P. Sinke, E. Swartz, H. Sanctorum, C. van der Giesen and I. Odegard, Ex-ante life cycle assessment of commercial-scale cultivated meat production in 2030, Int J Life Cycle Assess, 2023, 28, 234–254. 10.1007/s11367-022-02128-8

3 P. G. Negulescu, D. Risner, E. S. Spang, D. Sumner, D. Block, S. Nandi and K. A. McDonald, Techno-economic modeling and assessment of cultivated meat: Impact of production bioreactor scale, Biotechnology and Bioengineering, 2023, 120, 1055–1067. 10.1002/bit.28324.

4 C. Semper and A. Savchenko, Protein expression and purification of bioactive growth factors for use in cell culture and cellular agriculture, STAR Protoc, 2023, 4, 102351. 10.1016/j.xpro.2023.102351.

5 H.-H. Kuo, X. Gao, J.-M. DeKeyser, K. A. Fetterman, E. A. Pinheiro, C. J. Weddle, H. Fonoudi, M. V. Orman, M. Romero-Tejeda, M. Jouni, M. Blancard, T. Magdy, C. L. Epting, A. L. George and P. W. Burridge, Negligible-Cost and Weekend-Free Chemically Defined Human iPSC Culture, Stem Cell Reports, 2020, 14, 256–270. 10.1016/j.stemcr.2019.12.007.

6 A. J. Stout, A. B. Mirliani, M. L. Rittenberg, M. Shub, E. C. White, J. S. K. Yuen and D. L. Kaplan, Simple and effective serum-free medium for sustained expansion of bovine satellite cells for cell cultured meat, Commun Biol, 2022, 5, 466. 10.1038/s42003-022-03423-8.

7 L. Specht, The Good Food Institute, 2020. An analysis of culture medium costs and production volumes for cultivated meat, https://gfi.org/wp-content/uploads/2021/01/clean-meat-production-volume-and-medium-cost.pdf, (accessed January 2026).

8 K. R. Trinidad, R. Ashizawa, A. Nikkhah, C. Semper, C. Casolaro, D. L. Kaplan, A. Savchenko and N. T. Blackstone, Environmental life cycle assessment of recombinant growth factor production for cultivated meat applications, Journal of Cleaner Production, 2023, 419, 138153. 10.1016/j.jclepro.2023.138153.

9 M. Venkatesan, C. Semper, S. Skrivergaard, R. Di Leo, N. Mesa, M. K. Rasmussen, J. F. Young, M. Therkildsen, P. J. Stogios and A. Savchenko, Recombinant production of growth factors for application in cell culture, iScience, 2022, 25, 105054. 10.1016/j.isci.2022.105054

10 P. Mainali, M. S.-W. Chua, D.-J. Tan, B. L.-W. Loo and D. S.-W. Ow, Enhancing recombinant growth factor and serum protein production for cultivated meat manufacturing, Microb Cell Fact, 2025, 24, 41. 10.1186/s12934-025-02670-8

11 L. Li, B. Yu, Y. Lai, S. Shen, Y. Yan, G. Dong, X. Gao, Y. Cao, C. Ge, L. Zhu, H. Liu, S. Tao, Z. Yao, S. Li, X. Wang and Q. Hui, Scaling up production of recombinant human basic fibroblast growth factor in an Escherichia coli BL21(DE3) plysS strain and evaluation of its pro-wound healing efficacy, Front. Pharmacol. 2024, 14. 10.3389/fphar.2023.1279516.

12 D. G. Sauer, M. Mosor, A.-C. Frank, F. Weiß, A. Christler, N. Walch, A. Jungbauer and A. Dürauer, A two-step process for capture and purification of human basic fibroblast growth factor from E. coli homogenate: Yield versus endotoxin clearance, Protein Expr Purif, 2019, 153, 70–82. 10.1016/j.pep.2018.08.009.

13 C. D. Lynch and D. J. O’Connell, Conversion of mammalian cell culture media waste to microbial fermentation feed efficiently supports production of recombinant protein by Escherichia coli, PLoS One, 2022, 17, e0266921. 10.1371/journal.pone.0266921.

14 J. Rizal, P. Mainali, J. P. Quek, L. L. Tan, J. Bi, A. J. Chan, A. A. Gaffoor, L. J. M. Chew, S. Sugii, S. K. Ng, D. S.-W. Ow and F. T. Wong, Valorisation of spent cultivated meat media for recombinant FGF2 production in Lactococcus lactis, Syst Microbiol and Biomanuf, 2025, 5, 1328–1334. 10.1007/s43393-025-00358-z.

15 Q. Lei, J. Ma, G. Du, J. Zhou and X. Guan, Efficient expression of a cytokine combination in Saccharomyces cerevisiae for cultured meat production, Food Research International, 2023, 170, 113017. 10.1016/j.foodres.2023.113017.

16 J. Dolgin, D. Chakravarty, S. F. Sullivan, Y. Cai, T. Lim, P. Yamaguchi, J. E. Balkan, L. Xu, A. D. Olawoyin, K. Lee, D. L. Kaplan and N. U. Nair, Microbial lysates as low-cost serum replacements in cellular agriculture media formulation, Food Research International, 2025, 201, 115633. 10.1016/j.foodres.2024.115633.

17 A. M. Kolkmann, A. Van Essen, M. J. Post and P. Moutsatsou, Development of a Chemically Defined Medium for in vitro Expansion of Primary Bovine Satellite Cells, Front Bioeng Biotechnol, 2022, 10, 895289. 10.3389/fbioe.2022.895289.

18 Y. A. Seo, M. J. Cha, S. Park, S. Lee, Y. J. Lim, D. W. Son, E. J. Lee, P. Kim and S. Chang, Development of a Normal Porcine Cell Line Growing in a Heme-Supplemented, Serum-Free Condition for Cultured Meat, Int J Mol Sci, 2024, 25, 5824. 10.3390/ijms25115824.

19 N. Dong, B. Jiang, Y. Chang, Y. Wang and C. Xue, Integrated Omics Approach: Revealing the Mechanism of Auxenochlorella pyrenoidosa Protein Extract Replacing Fetal Bovine Serum for Fish Muscle Cell Culture, J. Agric. Food Chem., 2024, 72, 6064–6076. 10.1021/acs.jafc.4c00624.

20 Y. Okamoto, Y. Haraguchi, A. Yoshida, H. Takahashi, K. Yamanaka, N. Sawamura, T. Asahi and T. Shimizu, Proliferation and differentiation of primary bovine myoblasts using Chlorella vulgaris extract for sustainable production of cultured meat, Biotechnology Progress, 2022, 38, e3239. 10.1002/btpr.3239.

21 A. J. Stout, M. L. Rittenberg, M. Shub, M. K. Saad, A. B. Mirliani, J. Dolgin and D. L. Kaplan, A Beefy-R culture medium: replacing albumin with rapeseed protein isolates, Biomaterials, 2023, 296, 122092. 10.1016/j.biomaterials.2023.122092.

22 M. T. Weinstock, E. D. Hesek, C. M. Wilson and D. G. Gibson, Vibrio natriegens as a fast-growing host for molecular biology, Nat Methods, 2016, 13, 849–851. 10.1038/nmeth.3970.

23 M. Smith, J. S. Hernández, S. Messing, N. Ramakrishnan, B. Higgins, J. Mehalko, S. Perkins, V. E. Wall, C. Grose, P. H. Frank, J. Cregger, P. V. Le, A. Johnson, M. Sherekar, M. Pagonis, M. Drew, M. Hong, S. R. T. Widmeyer, J.-P. Denson, K. Snead, I. Poon, T. Waybright, A. Champagne, D. Esposito, J. Jones, T. Taylor and W. Gillette, Producing recombinant proteins in Vibrio natriegens, Microb Cell Fact, 2024, 23, 208. 10.1186/s12934-024-02455-5.

24 D. A. Specht, T. J. Sheppard, F. Kennedy, S. Li, G. Gadikota and B. Barstow, Efficient natural plasmid transformation of Vibrio natriegens enables zero-capital molecular biology, PNAS Nexus, 2024, 3, pgad444. 10.1093/pnasnexus/pgad444.

25 A. J. Stout, M. J. Arnett, K. Chai, T. Guo, L. Liao, A. B. Mirliani, M. L. Rittenberg, M. Shub, E. C. White, J. S. K. Jr. Yuen, X. Zhang and D. L. Kaplan, Immortalized Bovine Satellite Cells for Cultured Meat Applications, ACS Synth. Biol., 2023, 12, 1567–1573. 10.1021/acssynbio.3c00216.

26 A. J. Stout, X. Zhang, S. M. Letcher, M. L. Rittenberg, M. Shub, K. M. Chai, M. Kaul and D. L. Kaplan, Engineered autocrine signaling eliminates muscle cell FGF2 requirements for cultured meat production, Cell Reports Sustainability, 2024, 1, 100009. 10.1101/2023.04.17.537163.

27 F. Relaix, D. Rocancourt, A. Mansouri and M. Buckingham, A Pax3/Pax7-dependent population of skeletal muscle progenitor cells, Nature, 2005, 435, 948–953. 10.1038/nature03594.

28 Y. Capetanaki, D. J. Milner and G. Weitzer, Desmin in muscle formation and maintenance: knockouts and consequences, Cell Struct Funct, 1997, 22, 103–116. 10.1247/csf.22.103.

29 T. Otsuka, H.-M. Kan, P. Y. Mengsteab, B. Tyson and C. T. Laurencin, Fibroblast growth factor 8b (FGF-8b) enhances myogenesis and inhibits adipogenesis in rotator cuff muscle cell populations in vitro, Proceedings of the National Academy of Sciences, 2024, 121, e2314585121. 10.1073/pnas.2314585121.

30 A. Hevaganinge, C. M. Weber, A. Filatova, A. Musser, A. Neri, J. Conway, Y. Yuan, M. Cattaneo, A. M. Clyne and Y. Tao, Fast-Training Deep Learning Algorithm for Multiplex Quantification of Mammalian Bioproduction Metabolites via Contactless Short-Wave Infrared Hyperspectral Sensing, ACS Omega, 2023, 8, 14774–14783. 10.1021/acsomega.3c00861.

31 C. P. Long, J. E. Gonzalez, R. M. Cipolla and M. R. Antoniewicz, Metabolism of the fast-growing bacterium Vibrio natriegens elucidated by 13C metabolic flux analysis, Metab Eng, 2017, 44, 191–197. 10.1016/j.ymben.2017.10.008.

32 X. Sun, Y. Shang, B. Zhang, P. Guo, Y. Luo and H. Wu, Engineering of fast-growing Vibrio natriegens for biosynthesis of poly(3-hydroxybutyrate-co-lactate), Bioresour. Bioprocess., 2024, 11, 86. 10.1186/s40643-024-00801-4.

33 H. Gu, Y. Kong, D. Huang, Y. Wang, V. Raghavan and J. Wang, Scaling Cultured Meat: Challenges and Solutions for Affordable Mass Production, Compr Rev Food Sci Food Saf, 2025, 24, e70221. 10.1111/1541-4337.70221.

34 D. Risner, P. Negulescu, Y. Kim, C. Nguyen, J. B. Siegel and E. S. Spang, Environmental Impacts of Cultured Meat: A Cradle-to-Gate Life Cycle Assessment, ACS Food Sci. Technol., 2025, 5, 61–74. 10.1021/acsfoodscitech.4c00281.

35 K. Yamanaka, Y. Haraguchi, H. Takahashi, I. Kawashima and T. Shimizu, Development of serum-free and grain-derived-nutrient-free medium using microalga-derived nutrients and mammalian cell-secreted growth factors for sustainable cultured meat production, Sci Rep, 2023, 13, 498. 10.1038/s41598-023-27629-w.

36 S. Skrivergaard, J. F. Young, N. Sahebekhtiari, C. Semper, M. Venkatesan, A. Savchenko, P. J. Stogios, M. Therkildsen and M. K. Rasmussen, A simple and robust serum-free media for the proliferation of muscle cells, Food Research International, 2023, 172, 113194. 10.1016/j.foodres.2023.113194.

37 R. da G. Ferreira, A. R. Azzoni and S. Freitas, Techno-economic analysis of the industrial production of a low-cost enzyme using E. coli: the case of recombinant β-glucosidase, Biotechnology for Biofuels, 2018, 11, 81. 10.1186/s13068-018-1077-0.

38 F. A. G. S. Silva, S. Branco, F. Dourado, B. Neto and M. Gama, Life cycle assessment of bacterial cellulose and comparison to other cellulosic sources, Journal of Cleaner Production, 2025, 493, 144876. 10.1016/j.jclepro.2025.144876.

39 H. Fuchs, S. R. Ullrich and S. Hedrich, Vibrio natriegens as a superior host for the production of c-type cytochromes and difficult-to-express redox proteins, Sci Rep, 2024, 14, 6093. 10.1038/s41598-024-54097-7.

40 S. S. González, O. Ad, B. Shah, Z. Zhang, X. Zhang, A. Chatterjee and A. Schepartz, Genetic Code Expansion in the Engineered Organism Vmax X2: High Yield and Exceptional Fidelity, ACS Cent. Sci., 2021, 7, 1500–1507. 10.1021/acscentsci.1c00499.

41 H. H. Lee, N. Ostrov, B. G. Wong, M. A. Gold, A. S. Khalil and G. M. Church, Functional genomics of the rapidly replicating bacterium Vibrio natriegens by CRISPRi, Nat Microbiol, 2019, 4, 1105–1113. 10.1038/s41564-019-0423-8.

42 J. Menetrey, C. Kasemkijwattana, C. S. Day, P. Bosch, M. Vogt, F. H. Fu, M. S. Moreland and J. Huard, Growth factors improve muscle healing in vivo, J Bone Joint Surg Br, 2000, 82, 131–137. 10.1302/0301-620x.82b1.8954.

43 C. D. Lynch, F. Cerrone, K. E. O’Connor and D. J. O’Connell, Feeding secondary fermentations with mammalian and fungal culture waste streams increases productivity and resource efficiency, RSC Sustainability, 2024, 2, 1868–1882. 10.1039/D3SU00483J.

44 J. P. Quek, A. A. Gaffoor, Y. X. Tan, T. R. M. Tan, Y. F. Chua, D. S. Z. Leong, A. S. Ali and S. K. Ng, Exploring cost reduction strategies for serum free media development, npj Sci Food, 2024, 8, 107. 10.1038/s41538-024-00352-0.

45 E. N. O’Neill, J. C. Ansel, G. A. Kwong, M. E. Plastino, J. Nelson, K. Baar and D. E. Block, Spent media analysis suggests cultivated meat media will require species and cell type optimization, npj Sci Food, 2022, 6, 46. 10.1038/s41538-022-00157-z.

46 C. Lüchtrath, E. Forsten, R. Polis, M. Hoffmann, A. S. Genis, A.-L. Kuhn, M. Hövels, U. Deppenmeier, J. Magnus and J. Büchs, Small-scale fed-batch cultivations of Vibrio natriegens: overcoming challenges for early process development, Bioprocess Biosyst Eng, 2025, 48, 1007–1024. 10.1007/s00449-025-03159-9.

47 C. M. Goodwin, M. Jabeen, B. M. Rao and R. A. Shirwaiker, Cell culture media for cultivated meat: Review and perspectives on first principles design to drive cost-effective scale-up, Future Foods, 2026, 13, 100908. 10.1016/j.fufo.2026.100908.

48 L. Pasitka, G. Wissotsky, M. Ayyash, N. Yarza, G. Rosoff, R. Kaminker and Y. Nahmias, Empirical economic analysis shows cost-effective continuous manufacturing of cultivated chicken using animal-free medium, Nat Food, 2024, 5, 693–702. 10.1038/s43016-024-01022-w.

