## Supplemental Information for "Engineered *Vibrio natriegens* lysate can replace multiple components of cell culture media"

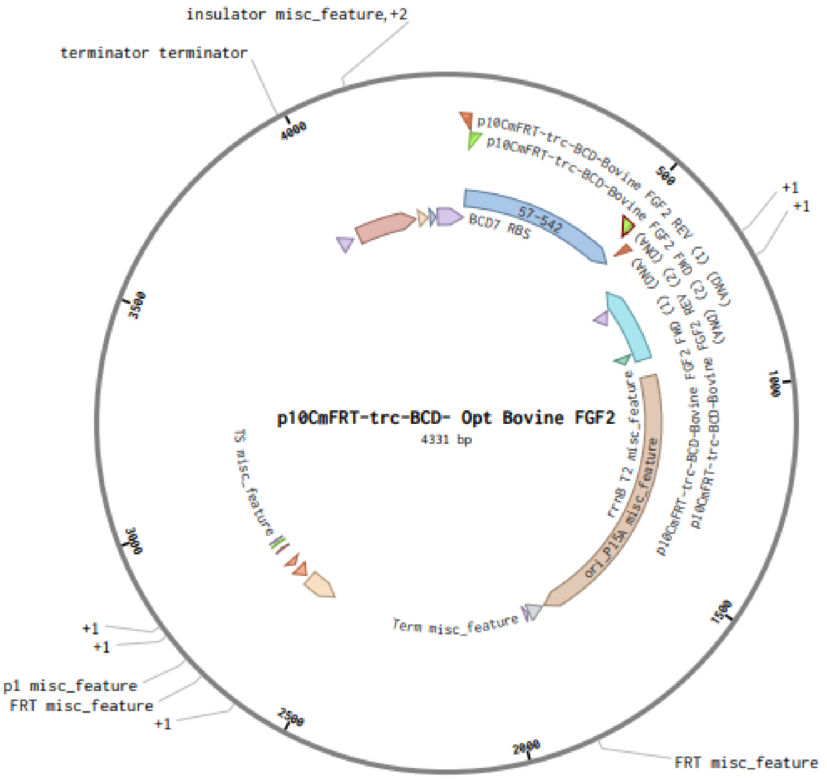


AcaatgaaagcaattttcgtactgaaacatcttaatcatgctggggagggtttctaATGGCGGC
AGGGAGTATTACTACGCTTCCCTCGTTACCAGAGGACGGAGGCTCTGGGGCCTTCCCACCAGGC
CACTTTAAGGACCCGAAGCGTCTTTACTGTAAAAATGGAGGGTTCTTCTTGCGTATTCACCCGG
ATGGGCGTGTTGACGGAGTACGCGAGAAGTCCGACCCTCATATCAAATTGCAGCTGCAGGCCGA
AGAACGCGGTGTAGTTTCCATTAAGGGAGTGTGTGCGAACCGCTATTTAGCAATGAAGGAAGAC
GGGCGTCTGCTGGCAAGCAAATGTGTCACGGACGAGTGTTTCTTTTTTGAGCGCCTTGAAAGCA
ATAATTACAACACCTATCGCTCGCGTAAGTATAGCAGTTGGTACGTTGCTCTTAAGCGTACAGG
CCAATACAAGCTGGGTCCCAAAACCGGCCCCGGCCAAAAAGCTATTCTTTTTTTGCCTATGAGC
GCAAAAAGTCACCACCACCACCACCACTGAtcgactagcgaattcgtcacgatcggtatcgagc
tctagccctaggagaagtgaaacgccgtagcgccgatggtagtgtggggtctccccatgcgaga
gtagggaactgccaggcatcaaataaaacgaaaggctcagtcgaaagactgggcctttcgtttt
atctgttgtttgtcggtgaacgctctcctgagtaggacaaatccgccgggagcggatttgaacg
ttgcgaagcaacggcccggagggtggcgggcaggacgcccgccataaactgccaggcatcaaat
taagcagaaggccatcctgacggatggcctttttcgcatgcagaacggccggtcagctagcgat
acctgcaggctgactgcagtagcaagcttgcgctagcggagtgtatactggcttactatgttgg
cactgatgagggtgtcagtgaagtgcttcatgtggcaggagaaaaaaggctgcaccggtgcgtc
agcagaatatgtgatacaggatatattccgcttcctcgctcactgactcgctacgctcggtcgt
tcgactgcggcgagcggaaatggcttacgaacggggcggagatttcctggaagatgccaggaag
atacttaacagggaagtgagagggccgcggcaaagccgtttttccataggctccgcccccctga
caagcatcacgaaatctgacgctcaaatcagtggtggcgaaacccgacaggactataaagatac
caggcgtttccccctggcggctccctcgtgcgctctcctgttcctgcctttcggtttaccggtgt
cattccgctgttatggccgcgtttgtctcattccacgcctgacactcagttccgggtaggcagt
tcgctccaagctggactgtatgcacgaaccccccgttcagtccgaccgctgcgccttatccggt
aactatcgtcttgagtccaacccggaaagacatgcaaaagcaccactggcagcagccactggta

```
attgatttagaggagttagtcttgaagtcatgcgccggttaaggctaaactgaaaggacaagtt
ttggtgactgcgctcctccaagccagttacctcggttcaaagagttggtagctcagagaacctt
cgaaaaaccgccctgcaaggcggttttttcgttttcagagcaagagattacgcgcagaccaaaa
cgatctcaagaagatcatcttattaatcagataaaatatttctagatttcagtgcaatttatct
cttcaaatgtagcacctgaagtcagccccatacgatgacgtcgaagttcctattccgaagttcc
tattctctagaaagtataggaacttcgactgccttaaaaaattacgccccgccctgccactca
tcgcagtactgttgtaattcattaagcattctgccgacatggaagccatcacagacggcatgat
gaacctgaatcgccagcggcatcagcaccttgtcgccttgcgtataatatttgcccatagtgaa
aacgggggcgaagaagttgtccatattggccacgtttaaatcaaaactggtgaaactcacccag
ggattggctgagacgaaaaacatattctcaataaaccctttagggaaataggccaggttttcac
cgtaacacgccacatcttgcgaatatatgtgtagaaactgccggaaatcgtcgtggtattcact
ccagagcgatgaaaacgtttcagtttgctcatggaaaacggtgtaacaagggtgaacactatcc
catatcaccagctcaccgtctttcattgccatacggaactccggatgagcattcatcaggcggg
caagaatgtgaataaaggccggataaaacttgtgcttatttttctttacggtctttaaaaaggc
cgtaatatccagctgaacggtctggttataggtacattgagcaactgactgaaatgcctcaaaa
tgttctttacgatgccattgggatatatcaacggtggtatatccagtgatttttttctccattt
tagcttccttagctcctgaaaatctcgataactcaaaaaatacgcccggtagtgatcttatttc
attatggtgaaagttggaacctcttacgtgccgatcaacggggatcctttgtgcgaagttcct
attctctagaaagtataggaacttccatatgctcgaggtaccaagcagctccagcctacaccaa
ttgcgcctgatcatgttcgtcatttacgttgacaccatcgaatggtgcaaaacctttcgcggta
tggcatgatagcgcccggaagagagtcaattcagggtggtgaatgtgaaaccagtaacgttata
cgatgtcgcagagtatgccggtgtctcttatcagaccgtttcccgcgtggtgaaccaggccagc
cacgtttctgcgaaaacgcgggaaaaagtggaagcggcgatggcggagctgaattacattccca
accgcgtggcacaacaactggcgggcaaacagtcgttgctgattggcgttgccacctccagtct
ggccctgcacgcgccgtcgcaaattgtcgcggcgattaaatctcgcgccgatcaactgggtgcc
agcgtggtggtgtcgatggtagaacgaagcggcgtcgaagcctgtaaagcggcggtgcacaatc
ttctcgcgcaacgcgtcagtgggctgatcattaactatccgctggatgaccaggatgccattgc
tgtggaagctgcctgcactaatgttccggcgttatttcttgatgtctctgaccagacacccatc
aacagtattattttctcccatgaagacggtacgcgactgggcgtggagcatctggtcgcattgg
gtcaccagcaaatcgcgctgttagcgggcccattaagttctgtctcggcgcgtctgcgtctggc
tggctggcataaatatctcactcgcaatcaaattcagccgatagcggaacgggaaggcgactgg
agtgccatgtccggttttcaacaaaccatgcaaatgctgaatgagggcatcgttcccactgcga
tgctggttgccaacgatcagatggcgctgggcgcaatgcgcgccattaccgagtccgggctgcg
cgttggtgcggatatctcggtagtgggatacgacgataccgaagacagctcatgttatatcccg
ccgtcaaccaccatcaaacaggattttcgcctgctggggcaaaccagcgtggaccgcttgctgc
aactctctcagggccaggcggtgaagggcaatcagctgttgcccgtctcactggtgaaaagaaa
aaccaccctggcgcccaatacgcaaaccgcctctccccgcgcgttggccgattcattaatgcag
ctggcacgacaggtttcccgactggaaagcgggcagtgatctgcgcgtaatctcttgctctgaa
aacgaaaaaaccgccttgcagggcggtttttcgaaggttctctgagctaccaactctttgaggc
aagccccctgaaattaatgccggcaggattacttttgatgaaacacagaggcgataatccggcc
ttcgcagcggccacttatactctgtgagccgaatccctggttgaaactgaaaatgcggcgctaa
actttcgtttcttatatagcaaaacttcatcaagcgcgctgatggataagcgcctcattaaccc
aaagatgaaatgagctgttgacaattaatcatccggctcgtataatgtgtggaattgtgagcgg
ataacaatttcacagggcccaagttcacttaaaaaggagatca...
```

**Bovine FGF2 gene in all caps

**Figure S1.** bFGF2 expression vector (plasmid map & sequence)

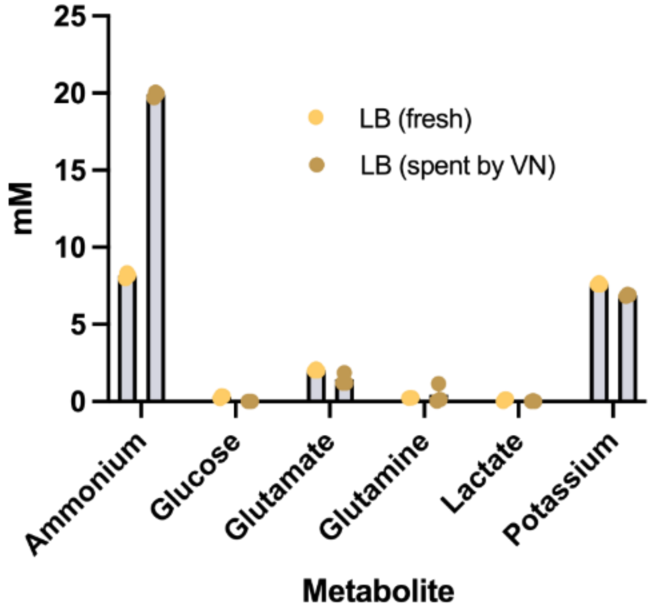


**Figure S2.** Metabolite analysis of fresh versus spent Luria Broth (LB)

**Table S1.** Cost breakdown of bovine satellite cell growth media

| BSCGM | | | | | | | | |
|---|---|---|---|---|---|---|---|---|
| **Component** | **Unit** | **Amount/ L** | **Supplier** | **Cat #** | **Units/ order** | **Cost $/order** | **$/Unit** | **$/L** |
| **DMEM+HEPES+Sodium bicarbonate** | mL | 800 | ThermoFisher | 11330057 | 5000 | 365 | 0.073 | 58.4 |
| **FBS** | mL | 200 | ThermoFisher | 26140079 | 500 | 660 | 1.32 | 264 |
| **FGF2** | μg | 1 | Peprotech | 100-18B | 1000 | 902 | 0.902 | 0.902 |
| **Total** | | | | | | | | **323.302** |

**Table S2.** Cost breakdown of serum-free Beefy-9 (B9) medium

| Component | Unit | Amount/L | Supplier | Cat# | Units/order | Unit cost ($) | $/Unit | $/L |
|---|---|---|---|---|---|---|---|---|
| **DMEM+HEPES+Sodium bicarbonate** | mL | 1000 | ThermoFisher | 11330057 | 5000 | 401 | 0.0802 | 80.2 |
| **L-ascorbic acid 2-phosphate** | µg | 200 | Sigma | 49752-250MG | 10000 | 85.8 | 0.00858 | 1.716 |
| **Insulin** | µg | 20 | Sigma | 91077C-250MG | 250 | 349 | 1.396 | 27.92 |
| **Transferrin** | µg | 20 | inVitria | 777TRI027-B | 1000 | 332.75 | 0.33275 | 6.655 |
| **Sodium selenite** | ng | 20 | Sigma | S5261-10G | 10000000 | 40.3 | 4.03E-6 | 8.1E-05 |
| **FGF2** | ng | 40 | Peprotech | 100-18B | 1000 | 902 | 0.902 | 36.08 |
| **NRG1** | ng | 100 | Peprotech | 396-HB-050 | 10000 | 89.5 | 0.00895 | 0.895 |
| **TGFβ3** | ng | 100 | R&D Systems | 243-B3-002 | 5000 | 249 | 0.0498 | 4.98 |
| **rAlbumin** | mg | 800 | Sigma | A9731-1G | 1000 | 133 | 0.133 | 106.4 |
| **Total** | | | | | | | | **264.846** |

**Table S3.** Cost breakdown of VN40 medium.

| Component | Unit | Amount/L | Supplier | Cat# | Units/order | Unit cost ($) | $/Unit | $/L |
|---|---|---|---|---|---|---|---|---|
| **DMEM+HEPES+Sodium bicarbonate** | mL | 1000 | ThermoFisher | 11330057 | 5000 | 401 | 0.0802 | 80.2 |
| **L-ascorbic acid 2-phosphate** | mg | 200 | Sigma | 49752-10G | 10000 | 85.8 | 0.00858 | 1.716 |
| **Insulin** | mg | 20 | Sigma | 91077C-250MG | 250 | 349 | 1.396 | 27.92 |
| **Transferrin** | mg | 20 | inVitria | 777TRF02 | 1000 | 332.75 | 0.33275 | 6.655 |
| **Sodium selenite** | µg | 20 | Sigma | S5261-10G | 10000000 | 40.3 | 4.03E-6 | 8.1E-05 |
| **FGF2** | µg | 40 | Peprotech | 100-18B | 1000 | 902 | 0.902 | 36.08 |
| **NRG1** | ng | 100 | Peprotech | 396-HB-050 | 10000 | 89.5 | 0.00895 | 0.895 |
| **TGFβ3** | ng | 100 | R&D Systems | s 8420-B3-005/CF | 5000 | 249 | 0.0498 | 4.98 |
| **VN Lysate** | mg | 40 | | | | | | 2 |
| **Total** | | | | | | | | **158.446** |

**Table S4.** Cost breakdown of VN40$^{FGF}$ medium.

| Component | Unit | Amount/L | Supplier | Cat# | Units/order | Unit cost ($) | $/Unit | $/L |
|---|---|---|---|---|---|---|---|---|
| **DMEM+HEPES+Sodium bicarbonate** | mL | 1000 | ThermoFisher | 11330057 | 5000 | 401 | 0.0802 | 80.2 |
| **L-ascorbic acid 2-phosphate** | mg | 200 | Sigma | 49752-10G | 10000 | 85.8 | 0.00858 | 1.716 |
| **Insulin** | mg | 20 | Sigma | 91077C-250MG | 250 | 349 | 1.396 | 27.92 |
| **Transferrin** | mg | 20 | inVitria | 777TRF02 | 1000 | 332.75 | 0.33275 | 6.655 |
| **Sodium selenite** | μg | 20 | Sigma | S5261-10G | 10000000 | 40.3 | 4.03E-6 | 8.1E-05 |
| **NRG1** | ng | 100 | Peprotech | 396-HB-050 | 10000 | 89.5 | 0.00895 | 0.895 |
| **TGFβ3** | ng | 100 | R&D Systems | s 8420-B3-005/CF | 5000 | 249 | 0.0498 | 4.98 |
| **VN Lysate** | mg | 40 | | | | | | 2 |
| **Total** | | | | | | | | **124.366** |

**Table S5.** Cost breakdown of *V. natriegens* lysate production using LB medium.

| Component | Unit | Amount/L | Supplier | Cat# | Units/order | Unit cost ($) | $/L LB | $/L VN40 |
|---|---|---|---|---|---|---|---|---|
| **Luria Broth** | g | 25 | ThermoFisher | 12795027 | 250 | 88 | 8.8 | 3.67 |
| **NaCl** | kg | 0.02 | Fisher Chemical | S271-500 | 1 | 100 | 2 | 0.83 |
| **Total** | | | | | | | | **4.50** |

**Table S6.** Cost breakdown of *V. natriegens* lysate production using spent culture medium.

| Component | Unit | Amount/L | Supplier | Cat# | Units/order | Unit cost ($) | $/L LB | $/L VN40 |
|---|---|---|---|---|---|---|---|---|
| **NaCl** | kg | 0.03 | Fisher Chemical | S271-500 | 1 | 100 | 3 | 1.25 |
| **Total** | | | | | | | | **1.25** |
